# Using pan RNA-seq analysis to reveal the ubiquitous existence of 5’ end and 3’ end small RNAs

**DOI:** 10.1101/444117

**Authors:** Xiaofeng Xu, Haishuo Ji, Zhi Cheng, Xiufeng Jin, Xue Yao, Yanqiang Liu, Qiang Zhao, Tao Zhang, Jishou Ruan, Wenjun Bu, Ze Chen, Shan Gao

## Abstract

In this study, we used pan RNA-seq analysis to reveal the ubiquitous existence of 5’ end and 3’ end small RNAs. 5’ and 3’ sRNAs alone can be used to annotate mitochondrial with 1-bp resolution and nuclear non-coding genes and identify new steady-state RNAs, which are usually from functional genes. Using 5’, 3’ and intronic sRNAs, we revealed that the enzymatic dsRNA cleavage and RNAi could involve in the RNA degradation and gene expression regulation of U1 snRNA in human. The further study of 5’, 3’ and intronic sRNAs help rediscover double-stranded RNA (dsRNA) cleavage, RNA interference (RNAi) and the regulation of gene expression, which challenges the classical theories. In this study, we provided a simple and cost effective way for the annotation of mitochondrial and nuclear non-coding genes and the identification of new steady-state RNAs, particularly long non-coding RNAs (lncRNAs). We also provided a different point of view for cancer and virus, based on the new discoveries of dsRNA cleavage, RNAi and the regulation of gene expression.

## Introduction

RNA sequencing (RNA-seq), usually based on the Next Generation Sequencing (NGS) technologies is widely used to simultaneously measure the expression levels of genes with higher accuracy than Serial Analysis of Gene Expression (SAGE) and microarray [1]. RNA-seq is also used to annotate genes in sequenced genomes, which is the only basis to study gene transcription, RNA processing and biological functions of these genes, *etc*. Particularly, RNA-seq or small RNA sequencing (sRNA-seq) is indispensable for the annotation of non-coding genes, while the annotation of protein-coding genes can be conducted based on the analysis of protein codons. However, RNA-seq cannot be used to obtain the full-length transcripts by *de novo* assembly or alignment. Both of PacBio full-length transcripts (PacBio cDNA-seq) [2] and Nanopore cDNA sequencing (Nanopore cDNA-seq) [1] can be used to obtain the full-length transcripts of mature RNAs or RNA precursors [3]. PacBio cDNA-seq produces reads with lower error rates than Nanopore cDNA-seq, while Nanopore cDNA-seq can produce longer reads than PacBio cDNA-seq. However, neither PacBio cDNA-seq nor Nanopore cDNA-seq can provide the exact 3’-end information of transcripts (*e.g.* polyA length) due to reverse transcription. The reason is that primers anneal to random positions located in the polyA regions or A-enriched regions in the body of transcripts to start reverse transcription. Nanopore direct RNA sequencing (Nanopore RNA-seq), as the only available sequencing technology which can sequence RNA directly [4], theoretically can be used to obtain the full-length 3’ ends of transcripts. But it can not be used to obtain the exact 3’-end information of transcripts either, due to the high error rate of Nanopore RNA-seq data. Combined with specific capture or enrichment technologies, several other RNA-seq methods have been developed to extent the use of standard RNA-seq. Parallel Analysis of RNA Ends and sequencing (PARE-seq), Cap Analysis of Gene Expression and sequencing (CAGE-seq) and Precision nuclear Run-On and sequencing (PRO-seq) have been developed to identify 5’ ends of mature RNAs. Polyadenylation sequencing (PA-seq) has been developed to identify 3’ ends of mature RNAs. Global Run-On and sequencing (GRO-seq) has been developed to sequence nascent RNAs [5], which helps determining the primary transcripts of genes.

The standard RNA-seq, sRNA-seq, PARE-seq, CAGE-seq, PRO-seq, PA-seq, GRO-seq, PacBio cDNA-seq, Nanopore cDNA-seq, Nanopore RNA-seq and *et al.*, defined as pan RNA-seq, were used to improve gene annotation in our previous studies. Using pan RNA-seq analysis, we reported the corrected annotation of tick rRNA genes, human rRNA genes [6], insect mitochondrial genes [3] and human mitochondrial genes [7]. We also reported two novel long non-coding RNAs (lncRNAs) discovered in human mitochondrial DNA [7]. In addition, we unexpectedly discovered the existence of 5’ and 3’ end small RNAs (5’ and 3’ sRNAs) in animal rRNA genes [6] and later proved the ubiquitous existence of 5’ and 3’ sRNAs in mitochondrial and nuclear non-coding genes. In this study, we demonstrated that 5’ and 3’ sRNAs alone can be used to annotate mitochondrial and nuclear non-coding genes with 1-bp resolution and identify new steady-state RNAs. Using public sRNA-seq data from the same species, this method provides a simple and cost effective way for the annotation of mitochondrial and nuclear non-coding genes and the identification of new steady-state RNAs, which are usually from functional genes. The further study of 5’, 3’ and intronic sRNAs help rediscover double-stranded RNA (dsRNA) cleavage, RNA interference (RNAi) and the regulation of gene expression, which challenges the classical theories.

## Results

### Discovery of 5’ and 3’ sRNAs

In a genome-alignment map of sRNA data, there usually are some peaks or hotspots [8], where the depths of the positions are much higher than those of other positions in the genome. In our previous study of human rRNA genes [6], we found that some of peaks comprised 5’ and 3’ sRNAs and they were ubiquitously existed in mitochondrial and nuclear non-coding genes. As the current sRNA-seq technologies usually provide sequences with short lengths, 5’ and 3’ sRNAs are defined as sRNA-seq reads with lengths of 15~50 bp, which are precisely aligned to the 5’ and 3’ ends of mature RNAs respectively (Figure 1A) and they have such features: **1)** 5’ and 3’ sRNAs are degraded fragments from mature RNAs and the lengths of them vary progressively with 1-bp differences. **2)** The cleavage sites between 3’ sRNAs and their downstream 5’ sRNAs are not limited to one (usually three) due to inexact cleavage by enzymes (Figure 1B). **3)** 5’ and 3’ sRNAs of steady-state RNAs (*e.g.* 18S, 5.8S and 28S rRNA) are significantly more abundant than the intronic sRNAs of them, while 5’ and 3’ sRNAs of transicent RNAs (*e.g.* Internal Transcribed Spacers of rRNA, ITS1 and ITS2) are not. This criterion can be used to identify new steady-state RNAs, which are usually from functional genes. One example of a new steady-state RNA downstream tRNAs and another example of two novel mitochondrial lncRNAs were introduced in the following paragraphs. Particularly, it was proved that MDL1 and MDL1AS were two steady-state lncRNAs in the human mitochondrial DNA and predicted to have biological functions [7].

**Figure 1.**
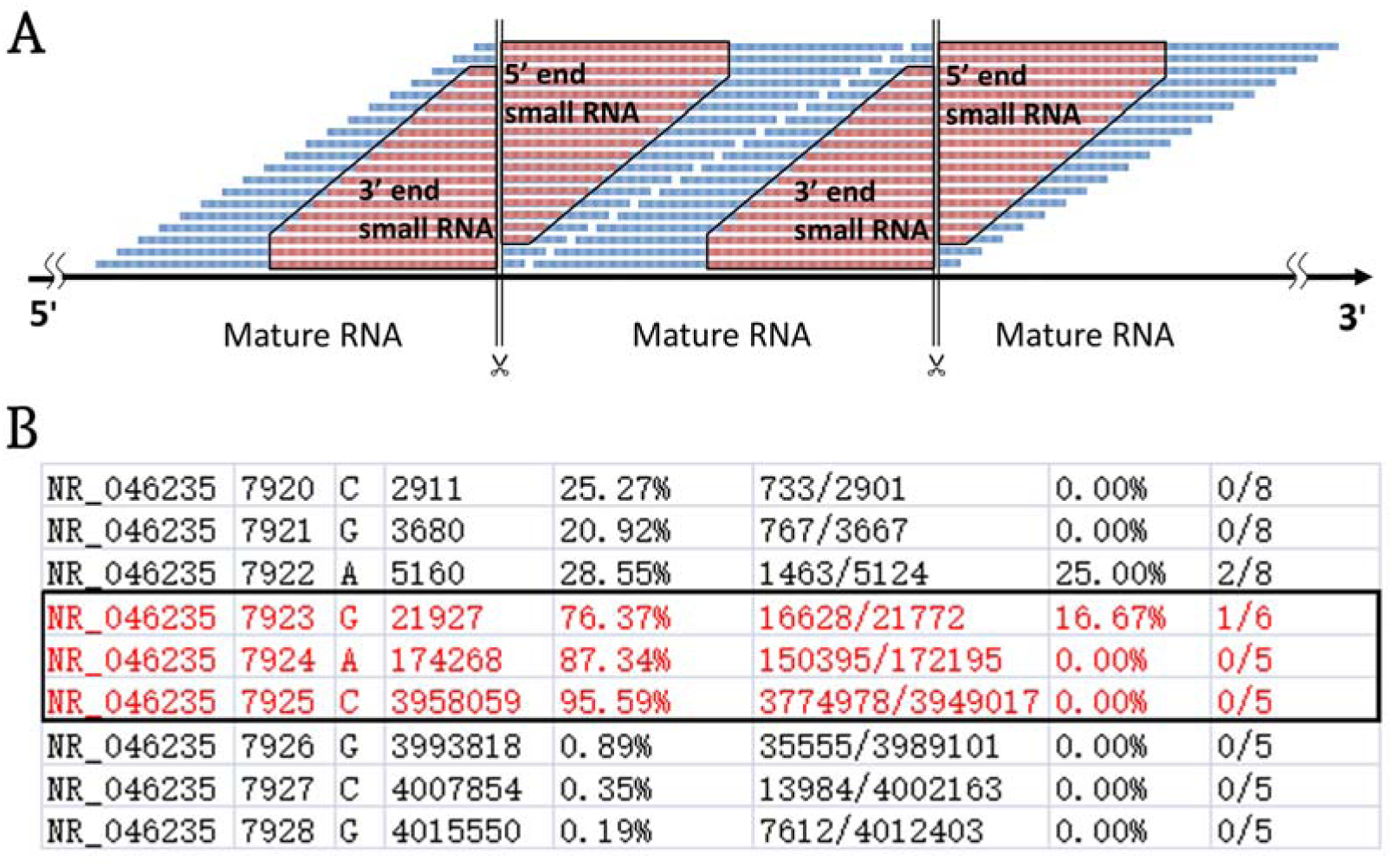
Definition of 5’ and 3’ sRNAs. **A.** 5’ and 3’ sRNAs are defined as sRNA-seq reads with lengths of 15~50 bp, which are precisely aligned to the 5’ and 3’ ends of mature RNAs respectively. The lengths of them vary progressively with 1-bp differences. **B.** 5-end format is defined to easily identify 5’ ends of mature RNAs using sRNA-seq data. Human rRNA genes (RefSeq: NR_046235.1) were annotated using alignment results in the 5-end format. Among positions 7923, 7924 and 7925 with ratio1s (the 5^th^ column) above 70 %, 7925 with the highest ratio1 was determined as the 5’ end of 28S rRNA.

We used 5’ and 3’ sRNAs from one sRNA-seq dataset to annotate genes and used one CAGE-seq dataset, one GRO-seq dataset and one PacBio cDNA-seq dataset (**Materials and Methods**) to validate the annotations. Later, we developed a simplified gene-annotation procedure. Using only 5’ sRNAs, gene annotation can be reduced to the identification of 5’ ends of mature RNAs. By doing so, the 3’ ends of their upstream mature RNAs and their cleavage sites can be derived (Figure 1A). We have defined a new file format, named 5-end format, to easily identify 5’ ends of mature RNAs. The new format is derived from the Pileup format (**Materials and Methods**) to include eight columns (Figure 1B) for each line including information from a genomic position: **1)** chromosome ID, **2)** 1-based coordinate of this position, **3)** reference base, **4)** depth (the number of reads covering the position), **5)** ratio1 (the number of positive-stranded reads starting at this position divided by the number of positive-stranded reads), **6)** the number of positive-stranded reads starting at this position divided by the number of positive-stranded reads, **7)** ratio2 (the number of negative-stranded reads starting at this position divided by the number of negative-stranded reads), **8)** the number of negative-stranded reads starting at this position divided by the number of negative-stranded reads. Using the 5-end format, the 5’ end of one mature RNA can be easily identified from two to three candidates (Figure 1B). Their ratio1s or ratio2s must be above a threshold (*e.g.* 75%) and significantly higher than the ratio1s or ratio2s of positions surrounding these two to three positions.

### 5’ and 3’ sRNAs in non-coding genes

Using 5’ and 3’ sRNAs, we corrected the annotation of human rRNA genes. For the 5’ end of each mature RNA, we obtained two or three candidates and selected the position with the highest ratio1 or ratio2 as the result. For example (Figure 1B), we obtained three positions 7,923, 7,924 and 7,925 to identify the 5’ end of 28S rRNA and selected 7,925 as the result. In the same way, 5’ ends of 18S and 5.8S rRNA were also identified using 5’ sRNAs. Then, 3’ ends of 18S, 5.8S and 28S rRNA were identified using 3’ sRNAs. Finally, the annotations of ITS1 and ITS2 were derived using the annotations of 18S, 5.8S and 28S rRNA. These corrected annotations (**Table 2**) were validated using the CAGE-seq dataset and the GRO-seq dataset (**Materials and Methods**). Although the depth 1,471,247 at the position 6,601 was much higher than the depth 647,406 at the position 6,596 in the sRNA-seq dataset, the 5’ end of 5.8S rRNA annotated as the position 6,601 with the ratio1 of 35.42% (520,006/1,468,024) was still corrected as the position 6,596 with the ratio1 of 88.11% (569,882/646,805). In addition, the genome-alignment map using the sRNA-seq dataset showed that human rRNA genes had peaks at the position 6,596, 7,925 and 6,756 corresponding to the 5’ ends of 5.8S and 28S rRNA and the 3’ end of 5.8S rRNA, respectively (Figure 2A). The genome-alignment map using the CAGE-seq dataset showed that human rRNA genes had peaks at the position 3,675 and 7,926 corresponding to the 5’ ends of 18S and 28S rRNA, respectively (Figure 2B). This suggested that 5’ m^7^G or other caps of 18S and 28S rRNA could exist. By the analysis of 3’ sRNAs, we confirmed that rRNA genes did not contain 3’ polyAs.

**Figure 2.**
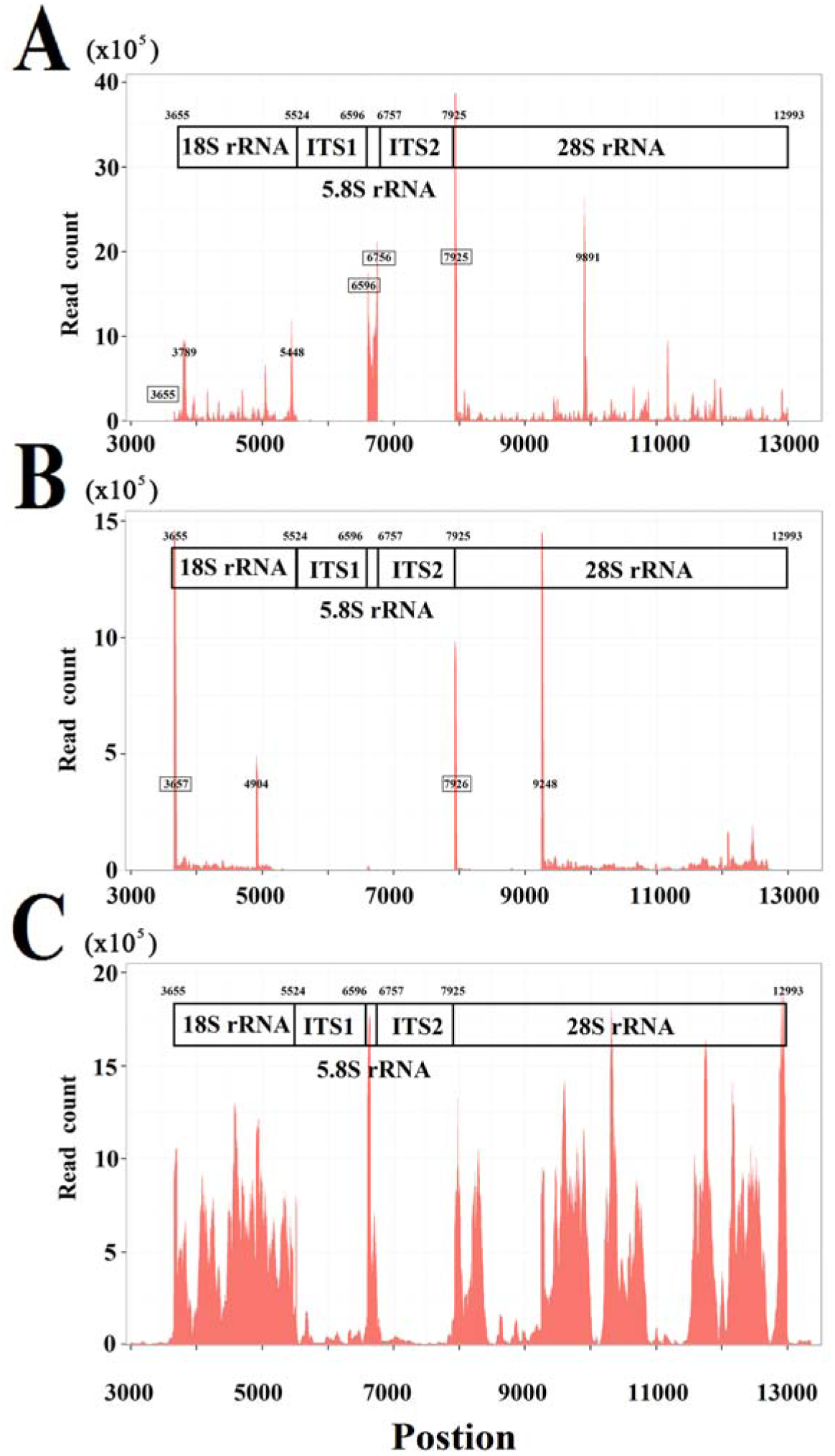
Genome-alignment maps using sRNA-seq, Cage-seq and GRO-seq. This figure shows the count distribution of all aligned reads on the reference rRNA sequence (RefSeq: NR_046235.1). These reads are from one sRNA-seq dataset (**A.**), one CAGE-seq dataset (**B.**) and one GRO-seq dataset (**C.**). The description of these datasets can be seen in the section Materials and Methods. The identified 5’ and 3’ ends of mature RNAs are marked by boxes.

In 2009, a novel class of sRNAs named tRNA-derived RNA fragments (tRFs) was introduced and three series of tRFs (tRF-5, tRF-3 and tRF-1) were identified using the sRNA-seq data of the human prostate cancer cell line by 454 deep sequencing [9]. However, these authors did not correctly understand the tRFs due to the technology limit and the small dataset size. Using the pan RNA-seq analysis, we proved that the tRF-5 and tRF-3 series were 5’ and 3’ sRNAs from mature tRNAs and the tRF-1 series were 5’ sRNAs from mature RNAs of their downstream genes (Figure 2A). In this study, 13 mature tRNAs and their 43 precursors (**Supplementary file 1**) were analyzed, and the 5/3-sRNA ratios of them ranged from 0.1 to 595.8. However, we did not found the characteristics in the size distribution of 5’ and 3’ sRNAs from these tRNAs. As 3’ sRNAs contain detailed 3’-end information of mature RNAs, we acquired more discoveries related to tRNA processing, maturation and degradation. One example was 3’ sRNAs of tRNAs had four types, which were non-tail, C-, CC-, and CCA-tailed. The proportions of these four types were 5.26% (22,906/4,355,95), 12.36% (53,845/4,355,95), 13.81% (60,176/4,355,95) and 68.57% (298,668/4,355,95). In addition, we obtained sequences of full-length mature tRNAs with non-tail, C-, CC-, and CCA-tailed. Among these full-length mature tRNAs, 8,539 TRD-GTC2-1 tRNAs (for Asp) and 16,900 TRE-CTC1-1 tRNAs (for Glu) were obtained. These results suggested that 3’ sRNAs were produced by tRNA degradation during its synthesis, when CCAs were post-transcriptionally added to 3’ ends of tRNAs one nucleotide by one nucleotide. Another example was the correction of TRL-TAG3-1’s annotation. The mature TRL-TAG3-1 was annotated as a 82-nt sequence from the human genome with its 3’ cleavage site ACCGCTGCCA|cacctcagaa. Using 5’ and 3’ sRNAs, the 3’ cleavage site of TRL-TAG3-1 was determined as ACCGCTGCCAC|acctcagaa. Instead of CAA, it was CA that was post-transcriptionally added to the 3’ end of TRL-TAG3-1 ACCGCTGCCAC. The genome-alignment results using the CAGE-seq dataset showed that 5’ m^7^G or other caps of tRNA did not exist. By the analysis of 3’ sRNAs, we confirmed that tRNA genes did not contain 3’ polyAs. 5’ and 3’ sRNAs from all the 13 mature tRNAs resided in peaks in the genome-alignment maps, while a few 3’ sRNAs of their upstream genes or 5’ sRNAs of their downstream genes resided in peaks. Among the peaks from these upstream or downstream genes, the highest one was on the downstream of TRS-TGA1-1 (chr10:67764503-67764584). It suggested that this peak was the 5’ end of a new steady-state RNA, which could be from a functional gene and had not been annotated in the current genome version.

Small nuclear RNAs (snRNAs) include a class of small RNA molecules that are found within the splicing speckles and Cajal bodies of the cell nucleus in eukaryotic cells [10]. snRNAs are always associated with a set of specific proteins and the complexes are referred to as small nuclear ribonucleoproteins (snRNPs). SnRNAs are also commonly referred to as U-RNAs and one well-known member is U1 snRNA [11]. Using 5’ sRNAs, we confirmed annotations of U1, U2, U3, U4, U5, U6 and U7 (**Supplementary file 1**). The genome-alignment results using the CAGE-seq dataset showed that U1, U2, U3 and U4 snRNAs could be capped by 5’ m^7^G, but U5, U6 and U7 snRNAs could not. By the analysis of 3’ sRNAs, we confirmed that all the snRNAs did not contain 3’ polyAs. In addition, we did not found any new steady-state RNA on the upstream or downstream of seven snRNA genes.

### 5’ and 3’ sRNAs in mitochondrial genes

Using pan RNA-seq analysis, we confirmed that nuclear mitochondrial DNA segments (NUMTs) in human genome did not transcribe into RNAs [7]. This finding simplified the analysis of mitochondrial genes (*e.g.* mutation detection or quantification) using RNA-seq data. In our previous study, we annotated two primary transcripts and 30 mature transcripts (tRNA^Ile^, tRNA^Gln^AS, tRNA^Met^, ND2, tRNA^Trp^, tRNA^Ala^AS/tRNA^Asn^AS/tRNA^Cys^AS/tRNA^Tyr^AS, COI, tRNA^Ser^AS, tRNA^Asp^, COII, tRNA^Lys^, ATP8/6, COIII, tRNA^Gly^, ND3, tRNA^Arg^, ND4L/4, tRNA^His^, tRNA^Ser^, tRNA^Leu^, ND5/ND6AS/tRNA^Glu^AS, Cytb, tRNA^Thr^, MDL1, tRNA^Phe^, 12S rRNA, tRNA^Val^, 16S rRNA, tRNA^Leu^ and ND1) on the H-strand with 1-bp resolution [7]. In this study, COI and tRNA^Ser^AS were corrected as one mature transcript COI/tRNA^Ser^AS that could not be further cleaved. We classified mitochondrial genes into tRNA, mRNA, rRNA, antisense tRNA (*e.g.* tRNA^Ser^AS), antisense mRNA (*e.g.* ND6AS), antisense rRNA and lncRNAs (*e.g.* MDL1 and MDL1AS) [7] and improved the "mitochondrial cleavage" model that had been proposed in our previous study [7]. The improved model is that RNA cleavage can be processed: 1) at 5’ and 3’ ends of tRNAs, 2) between mRNAs and mRNAs (*e.g.* ATP8/6 and COIII) and 3) between antisense tRNAs and mRNAs (*e.g.* tRNA^Tyr^AS and COI), but cannot be processed: 1) between mRNAs and antisense tRNAs (*e.g.* COI and tRNA^Ser^AS) 2) between mRNAs and antisense mRNAs (*e.g.* ND5 and ND6AS), 3) between antisense mRNAs and antisense tRNAs (*e.g.* ND6AS and tRNA^Glu^AS) or 4) between antisense tRNAs and antisense tRNAs (*e.g.* tRNA^Ala^AS/tRNA^Asn^AS/tRNA^Cys^AS/tRNA^Tyr^AS). This model provided a framework to identify all the full-length coding and non-coding RNAs of animal mitochondrion with 1-bp resolution. For example, the identification of ND5/ND6AS/tRNA^Glu^AS, MDL1 and MDL1AS proved that all the reported lncRNAs from ND5, ND6, Cytb [12] or the D-loop region were meaningless fragments from RNA degradation or. Another example was that tRNA^Ala^AS-tRNA^Tyr^AS (NC_012920: 1318-1638) could not be further cleaved, which was against a hypothesis from one previous study [13].

Among 29 mature transcripts on the H-strand, tRNA transcripts were tailed by 3’ CCAs, while other mature transcripts were tailed by 3’ polyAs. The maximum lengths of polyAs in tRNA^Gln^AS, ND2, tRNA^Ala^AS-tRNA^Tyr^AS, COI/tRNA^Ser^AS, COII, ATP8/6, COIII, ND3, ND4L/4, ND5/ND6AS/tRNA^Glu^AS, Cytb, MDL1, 12S rRNA, 16S rRNA, and ND1 are 14, 1, 12, 11, 20, 35, 10, 1, 22, 1, 8, 28, 32, 21 and 16, respectively. 3’ sRNAs containing polyAs or CCAs of different lengths were captured to proved that 3’ sRNAs were produced by RNA degradation during its synthesis, when polyAs or CCAs were post-transcriptionally added to 3’ ends of RNAs one nucleotide by one nucleotide. In this study, we also confirmed that both of mRNA and rRNA transcripts were capped by 5’ m^7^G [3]. Our data supported that MDL1AS, ND5/ND6AS/tRNA^Glu^AS and tRNA^Ala^AS/tRNA^Asn^AS/tRNA^Cys^AS/tRNA^Tyr^AS could be capped by 5’ m^7^G, but tRNA^Gln^AS and MDL1 could not be capped. 5’ and 3’ sRNAs of MDL1 and MDL1AS were significantly more abundant than the intronic sRNAs of them, which was one criterion to identify steady-state RNAs. Although MDL1 was not capped by 5’ m^7^G as MDL1AS, we still proposed that they were steady-state RNAs and could have biological functions. The further study showed that qPCR of MDL1 provided higher sensitivities than that of BAX/BCL2 and CASP3 in the detection of cell apoptosis [14].

In our previous study, the first Transcription Initiation Site (TIS) of H-strand (IT_H1_) and the TIS of L-strand (IT_L_) were determined at the position 561 and 407 on the human mitochondrial genome (RefSeq: NC_012920.1), but the second TIS of H-strand (IT_H2_) was not determined [7]. In this study, IT_H2_ was determined at the position 648, which was also the 5’ end of 12S rRNA. This finding was against the long-standing knowledge that IT_H2_ was at the position 638 [15]. Using pan RNA-seq analysis, we found that all the TISs (IT_H1,_ IT_H2_ and IT_L_) could be capped by 5’ m^7^G. We also found polyAs before TISs, which suggested that the transcription of mitochondrial genes could be initated by primers containing polyTs. This finding explained why all the TISs were resided in A-enriched regions. However, these explanations need be proved in the future studies.

### Analysis of RNA degradation using 5’, 3’ and intronic sRNAs

As 5’, 3’ and intronic sRNAs are accumulated RNA degradation intermediates, they can be used to investigate the RNA degradation [16], particularly for steady-state RNAs. The result using our data showed that in general, 5’ and 3’ sRNAs were more abundant than intronic sRNAs and short 5’ and 3’ sRNAs were more abundant than longer ones for tRNAs, rRNAs, snRNAs and mitochondrial RNAs. This suggested that these mature RNAs, particularly short RNAs (*e.g.* tRNAs), were mainly degraded by 3’ and 5’ exonucleases to accumulate 5’ and 3’ sRNAs. As for rRNAs and snRNAs, we found many peaks in the body of genes, which were even much higher than the peaks comprising 5’ or 3’ sRNAs in the genome-alignment map. In addition, the peaks comprising intronic sRNAs in rRNAs showed tissue specificities. The liver tissue (SRA: SRP002272) had specific peaks at the positions 12,891. The Plasma (SRA: SRP034590) had specific peaks at the positions 5,431, 9,891 and 11,158. The B-Cell and exosome (SRA: SRP046046) had specific peaks at the positions 3,789 and 9,891. The Platelets (SRA: SRP048290) had specific peaks at the positions 4,384 and 10,627. Further study of these tissue specificities was beyond the scope of this study. Then, we focused on the study of the secondary structures around these peaks in rRNAs and snRNAs and found that some of them involved in dsRNA regions. Particularly, we found a featured peak spanning a 43-bp region from 49 bp to 92 bp of U1 snRNA (Figure 3). In this region, 5’ ends of most intronic sRNAs were precisely aligned to 49 bp or 78 bp (Figure 3A). We also found that this region formed a hairpin in the secondary structure of U1 snRNA and produced a series of siRNA duplexes [17] with lengths from 15 bp to at least 25 bp (Figure 3C). The most abundant reads AGGGCGAGGCTTATC and TGTGCTGACCCCTGC formed a 15-bp duplex structure. The duplex ratio of AGGGCGAGGCTTATC against TGTGCTGACCCCTGC was 2.15 (34,078/15,825) and 99.97% (49,889/49,903) of these duplexes were sequenced from 14 samples of plasma (SRA: SRP034590). It suggested that this dsRNA region was cleaved by the ribonuclease III (RNase III) family [18] to produce these siRNA duplexes and could induced RNAi. Based on the findings in this study, our hypothesis is: 5’ and 3’ exonucleases are prevalent than endonuceases for the degradation of mature non-coding RNAs. So abundant 5’ and 3’ sRNAs were observed using sRNA-seq data. The longer mature RNAs have more and longer dsRNA regions (*e.g.* 15-bp long for U1) than short ones (*e.g.* 7-bp longest for tRNAs) to induce dsRNA cleavage to produce siRNA duplexes. Although the lengths of siRNA duplexes discovered in this study were only 15 bp, we still hypothesized that they could induce RNAi due to the unbalanced duplex ratio 2.15. As RNAi regulate the expression of these genes through a negative feed-back mechanism, we designed preliminary experiments to over-express U1 snRNA in the HEK293 (human), SY5Y (human) and PC-12 (rat) cell lines to prove our hypothesis. The basic idea was that if the negative feed-back mechanism existed, the expression level of U1 snRNA could decrease rather than be stable when the over-expression of it beyond a threshold. The experimental results showed that the expression level of U1 snRNA decreased after 4X, 9X and 6X dosage (**Materials and Methods**) in the HEK293 (human), SY5Y (human) and PC-12 (rat) cell lines, respectively (Figure 3D). Particularly, the results in the HEK293 cell line showed a significant effect caused by the negative feed-back mechanism. Therefore, RNAi could involve in the RNA degradation and gene expression regulation of U1 snRNA.

**Figure 3.**
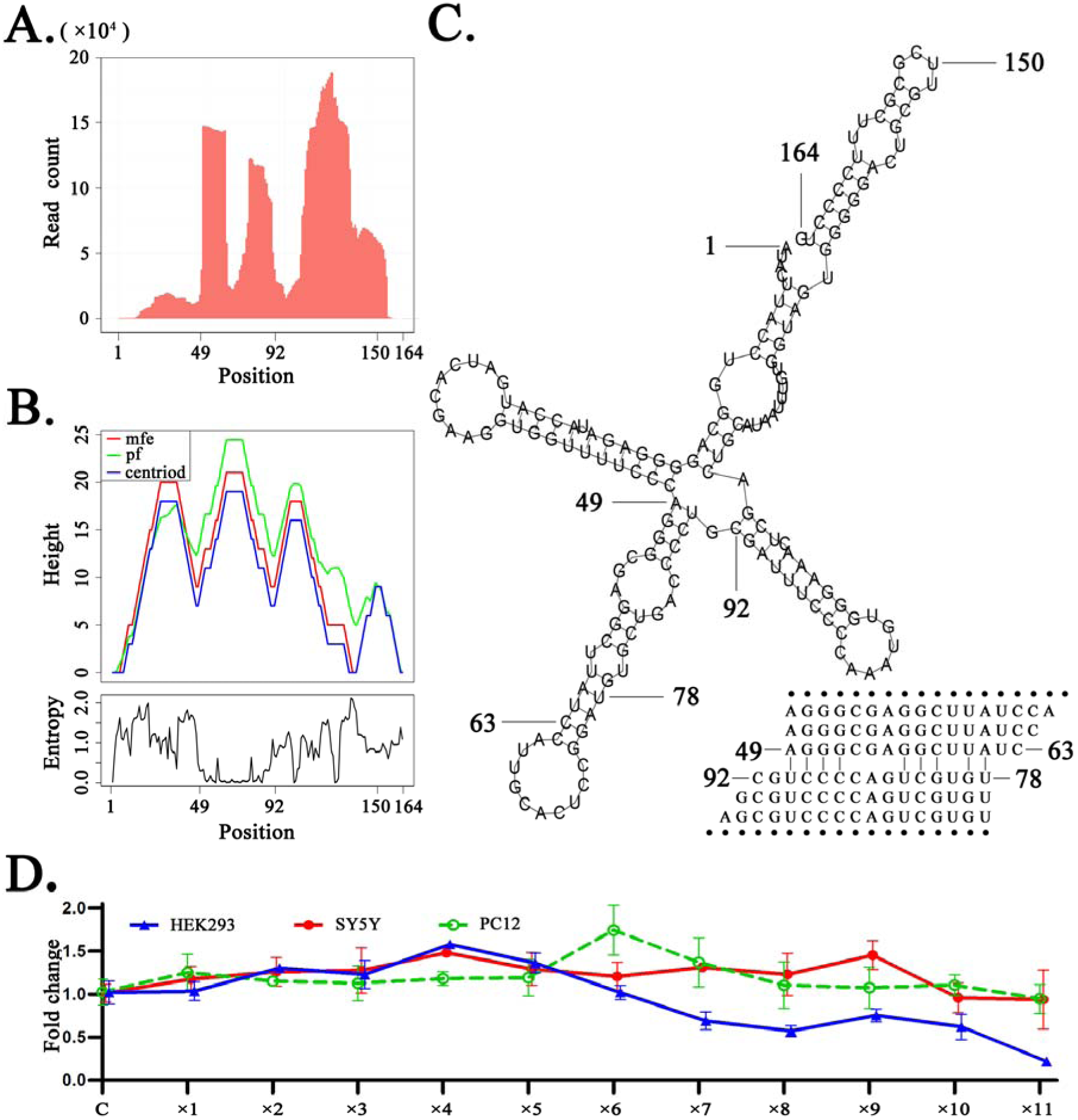
siRNA duplexes discovered from U1 snRNAs. **A.** The count distribution of all aligned reads on the reference U1 snRNA (RefSeq: NR_004430.2). **B.** The above is a mountain plot representation of the MFE structure, the thermodynamic ensemble of RNA structures and the centroid structure. The positional entropy for each position is showed below. **C.** The secondary structure of U1 snRNA. **D.** U1 over-expression in the HEK293 (human), SY5Y (human) and PC-12 (rat) cell lines were conducted by virus transfection. The qPCR results showed the relative expression levels of U1 in 12 groups (**Materials and Methods**). The control group used unprocessed samples.

## Conclusion and Discussion

In this study, we used the pan RNA-seq analysis to reveal the ubiquitous existence of 5’ end and 3’ end small RNAs. 5’ and 3’ sRNAs alone can be used to annotate mitochondrial and nuclear non-coding genes with 1-bp resolution and identify new steady-state RNAs. Using 5’, 3’ and intronic sRNAs, we revealed that the enzymatic dsRNA cleavage and RNAi could involve in the RNA degradation and gene expression regulation of U1 snRNA in human. The RNAi’s function in the RNA degradation was reported as a inappropriate event in yeast rRNA and tRNA degradation and only happened when 5’ and 3’ degradation were absent [19]. However, our finding suggested that RNAi’s function in the RNA degradation could be a general mechanism.

Based on a previous study, the Rnt1p polypeptide cleave hairpin structures in pre-rRNAs, pre-mRNAs, and transcripts containing noncoding RNAs (*e.g.* snoRNAs) for their maturation in yeast. Rnt1p recognizes the tetraloop [A/u]GNN and cleaves the stem ∼14–16 bp from the structure [18]. The most abundant read AGGGCGAGGCTTATC discovered in this study contained AGGG and AGGC tetraloops and had a length of 15 bp. It suggested that Rnt1p could produce those siRNA duplexes from U1 snRNAs and cause RNAi, which was against basic knowledge that Dicer is required for RNAi in mammal but produce siRNA duplexes with lengths of ∼20–25 bp. Although our preliminary experiments proved the existence of RNAi, which enzyme caused RNAi in U1 snRNAs is still unclear.

The ancestral function of RNAi is generally agreed to have been immune defense against exogenous genetic elements such as transposons and viral genomes [20]. However, our findings help rediscover dsRNA cleavage, RNAi and the regulation of gene expression. That is, both of dsRNA cleavage and RNAi are innate mechanisms rather than immune defense mechanisms. Basically, the enzymatic dsRNA cleavage is responsible for RNA processing, maturation and degradation, while RNAi is responsible for RNA degradation. By the degradation of mature RNAs, RNAi of one gene produces siRNA duplexes to regulate expression levels of itself or other genes. Mature RNAs containing more hairpin structures have more chances to induce RNAi, which is more important for highly expressed genes (*e.g.* U1 snRNA) or virus genes. These genes need RNAi to regulate gene expression though a negative feed-back mechanism. In one of our previous studies, we reported for the first time the existence of complemented palindromic small RNAs (cpsRNAs) and proposed that one cpsRNA from severe acute respiratory syndrome coronavirus (SARS-CoV) could induced RNAi [14]. This cpsRNA was detected from a 22-bp DNA complemented palindrome in the SARS-CoV genome. As DNA complemented palindromes are prone to produce dsRNA regions, viruses containing more DNA complemented palindromes in their genomes have more chances to induce RNAi and have more abilities for the regulation of gene expression, which is important for their infection or pathogenesis.

We also provided a different point of view for the gene expression regulation of U1 snRNA, The primary function of U1 snRNA is its involvement in the splicing of pre-mRNAs in nuclei. In the past 20 years, research of U1 snRNA has focused on its primary function, particularly related to neurodegenerative diseases caused by abnormalities of U1 snRNA [11]. In one of our previous studies, we reported that over-expression of U1 snRNA induces decrease of U1 spliceosome function associated with Alzheimer’s disease. However, the relationship between U1 snRNA over-expression and U1 snRNP loss of function remain unknown [21]. In another previous study, we reported that U1 snRNA over-expression induced cell apoptosis in SY5Y cells, but not in PC-12 cells [11]. These contradictory phenomena can be explained using the RNAi’s function in the RNA degradation and the negative feedback mechanism.

We provided a different point of view for cancer and virus. In one of our previous study, we reported how U1 snRNA over-expression affected the expression of mammal genes on a genome-wide scale and that U1 snRNA could regulate cancer gene expression. This had been explained by that Alternative Splicing (AS) and Alternative Polyadenylation (APA) were deregulated and exploited by cancer cells to promote their growth and survival [22]. Based on our point of view, the over-expressed U1 snRNA in cancer cells recruit excess RNase III. So the left RNase III is not enough to function in RNA degradation of other genes or genome surveillance and *etc* [18]. Viruses also recruit excess RNase III and the left RNase III is not enough to function in hose defense [18].

## Materials and Methods

### Datasets and data analysis

All sRNA-seq data were downloaded from the NCBI SRA database. Data in four projects (SRP002272, SRP034590, SRP046046 and SRP048290) were selected from the human931 sRNA-seq dataset which had been collected in our previous study [23]. SRP002272, SRP034590, SRP046046 and SRP048290 were sequenced using Illumina Single End (SE) sequencing technologies with length 35~46, 202, 101 and 101 bp, respectively and they contained 15, 14, 12 and 6 runs of sRNA-seq data, respectively. One CAGE-seq dataset, one GRO-seq dataset [5] and one PacBio cDNA-seq dataset [7] were used to validate the annotations (**Supplementary file 1)**. The cleaning and quality control of sRNA-seq data were performed using the pipeline Fastq_clean [24] that was optimized to clean the raw reads from Illumina platforms. To simply annotate genes from a sequenced genome, we aligned all the cleaned reads from sRNA-seq, CAGE-seq and GRO-seq data to the reference sequences using the software bowtie v0.12.7 allowing one mismatch. Then, we obtained SAM, BAM, sorted BAM, Pileup and 5-end files using the software samtools. Statistical computation and plotting were performed using the software R v2.15.3 with the Bioconductor packages [1].

### Validation by preliminary experiments

U1 over-expression in the HEK293 (human), SY5Y (human) and PC-12 (rat) cell lines were conducted by virus transfection using the pLVX-shRNA1 plasmids and the Lenti-X HTX Packaging System (Clontech, USA), which had been described in our previous study [21]. U1 snRNAs of human and rat used synthetic DNA containing the sequence (RefSeq: NR_004430.2) and the sequence (GenBank: V01266.1), respectively. For each experiment, 12 groups of samples named control, 1X, 2X, 3X, 4X, 5X, 6X, 7X, 8X, 9X, 10X and 11X were transfected by 0, 1, 2, 3, 4, 5, 6, 7, 8, 9, 10 and 11 μL U1-packged lentiviruses (Figure 3D). Each group contained three samples for biological replicates and the control samples used unprocessed cells. Each sample contained 10^5^ cells and virus titer was 10^7^ TU/mL for 1X. After transfection, RNA extraction, cDNA synthesis and cDNA amplification were performed following the same procedure in our previous study [11]. For each sample, total RNA was isolated using RNAiso Plus Reagent (TaKaRa, Japan) and the cDNA was synthesized by Mir-X miRNA First-Strand Synthesis Kit (Clontech, USA). The cDNA product was amplified by qPCR (Thermo Fisher Scientific, USA) using U6 snRNA as internal control under gene-specific reaction conditions. U1 snRNAs of human and rat used the forward and reverse primers GGGAGATACCATGATCAC and CCACTACCACAAATTATGC. U6 snRNAs of human and rat used CGGCAGCACATATACTAA and GAACGCTTCACGAATTTG.

**Table 1.** Annotation of human rRNA genes with corrections

| Gene | Start | End | Start* | End* | Length* |
| --- | --- | --- | --- | --- | --- |
| 18S rRNA | 3,655 | 5,523 | 3,655 | 5,523 | 1,869 |
| ITS1 | 5,524 | 6,600 | 5,524 | 6,595* | 1,072 |
| 5.8S rRNA | 6,601 | 6,757 | 6,596* | 6,756* | 161 |
| ITS2 | 6,758 | 7,924 | 6,757* | 7,924 | 1,168 |
| 28S rRNA | 7,925 | 12,994 | 7,925 | 12,993* | 5,069 |

Human rRNA genes (RefSeq: NR\_046235.1) were annotated using 5' and 3' sRNAs and \* represented the corrected annotation.

## Acknowledgments

We thank Yangguang Han and Jinke Huang from TianJin KeYiJiaXin Technology Co., Ltd for his professional guidance on the RNAi and cellular experiments.

## Authors’ contributions

SG conceived this project. SG and QW supervised this project. SG, ZC and ZW analyzed the data. YS and HC curated the sequences and prepared all the figures, tables and additional files. CL, WS and YH performed qPCR experiments. SG drafted the main manuscript. DY and WB revised the manuscript. **All authors have read and approved the manuscript.**

